# Growth inhibition in co-cultivation of *Cyclocybe aegerita* and *Hericium erinaceus* mushrooms

**DOI:** 10.1101/2025.10.23.684172

**Authors:** Carlos Wong, Robin A. Choudhury

## Abstract

Mushroom production relies heavily on compost and other substrates to grow and maintain fungal cultures. Large scale production of mushrooms can be achieved in specialized trays and grow rooms, however small-scale production of specialty mushrooms like lion’s mane (*Hericium erinaceus*) and pioppino mushrooms (*Cyclocybe aegerita*) relies on inoculation of small blocks of mushroom substrate in polypropylene bags, leaving spent mushroom substrate after production. Co-cultivation of mushrooms has been explored for the generation of novel secondary metabolites, however there is relatively little information about the effects of co-cultivation on yield. Our objective was to co-cultivate *H. erinaceus* and *C. aegerita* in the same bag and compare the overall colonization and yield with single inoculation bags. Overall, there was slower colonization and a significant reduction in the weight of harvested mushrooms from the mixed inoculation bag compared to the single inoculation bags, as well as a higher risk of contamination. We continue to explore new ways to improve the economic and environmental sustainability of mushroom production.

## 1. INTRODUCTION

Mushroom production in US is growing annually, with an estimated $1.09B in production in the 2023-2024 season (USDA NASS 2024). In addition to large-scale commercial production, many small-scale producers and hobbyists focus on specialty strains and varieties of mushrooms that cannot otherwise be bought commercially from supermarkets (Moxley et al. 2022). These small-scale producers rely on cheap and readily available substrates like soybean hulls and oak pellets for logistical and economic reasons and grow their mushrooms using mushroom bags (Fewell and Gustafson 2007; Haukongo 2021; Krupodorova and Barshteyn 2015). After these bags have been harvested, the spent mushroom substrate (SMS) is often discarded or used as mulch in agricultural settings, although there may be challenges associated with addition of SMS to container agriculture due to physical properties like high electrical conductivity (Catal and Peksen 2018). Efficiently using the substrate could help to improve yields and reduce waste streams.

Most mushrooms are produced by inoculating a single fungal isolate in the substrate to reduce risk of contamination, a major challenge for small producers (Moxley et al. 2022). However, there is a growing interest in co-cultivation of mushrooms for production of novel secondary metabolites not normally produced by single species of fungi (Carabajal et al. 2012; Knowles et al. 2022; Xu et al. 2023). Fungi use a complex set of enzymes to break down their preferred substrate, and many fungi are specialized in degradation of a specific type of substrate. Co-cultivation, growing multiple mushroom species in the same container, could offer a path towards higher mushroom yields and reduced waste. The objective of this experiment was to explore the effects of co-cultivation of two mushroom species (*Cyclocybe aegerita* and *Hericium erinaceus*) in a single bag compared to single inoculation bags to compare the colonization rate, contamination rate, and yield.

## 2. MATERIALS AND METHODS

Mushroom cultivation bags were prepared from 13L polypropylene bags with a 0.2-micron ventilation window that contained 0.9kg oak pellet and 0.9kg soybean hulls and hydrated with 2 liters of water per bag. Bags were sterilized and single-isolate grain spawn (North Spore LLC, Portland, ME) of *C. aegerita* and *H. erinaceus* and a mixture of the two isolates were inoculated into bags in a sterile flow hood. Six bags of each treatment were used for this experiment for a total of eighteen bags. The bags were maintained in grow tents that included a humidifier, a fan, and a UV light, allowing the mushrooms to have adequate environmental conditions to fruit and reproduce. Bags were monitored every 5-7 days for hyphal colonization and pinning as well as any visible contamination. Hyphal colonization was rated as a percent of visible substrate in the bag colonized by hyphae. Contamination was rated on an ordinal scale based on the severity and prevalence of visible contamination of substrate in the bag. Contamination was assessed by discolored hyphae and spores visible on the surface of the substrate. Twenty-eight days after inoculating the bags, the mushrooms from the bags were harvested and weighed. An ANOVA was run comparing the colonization of the mushroom bags for the three treatments using a Tukey’s Honestly Significant Difference test with an alpha of 0.05. Contamination was compared pairwise between the single isolate and co-cultivation bags using a ordinal logistic regression. Yield data was compared using a Wilcoxon rank sum exact test, comparing the yields collected from the species when grown singly or in co-cultivation. All statistics were conducted in R version 4.5.1 (R Core Team 2025).

## 3. RESULTS

Overall, *C. aegerita* had slower colonization of substrate when compared with *H. erinaceus*, with all six bags of *H. erinaceus* achieving >95% colonization within the first fifteen days, while *C. aegerita* did not achieve a similar level of colonization until day 28 (Fig. 1). There was a significant difference in hyphal colonization between all treatments on measurement day 5 (DF = 2, F = 87.7, p = 5.3e-9), day 12 (DF = 2, F = 37.3, p = 1.5e-6), and day 15 (DF = 2, F = 181, p = 3.2e-11), but no significant difference on day 20 (DF = 2, F = 2.4, p = 0.12), day 25 (DF = 2, F = 1.1, p=0.38), or day 28 (DF = 2, F = 1.02, p = 0.38).

**Figure 1.**
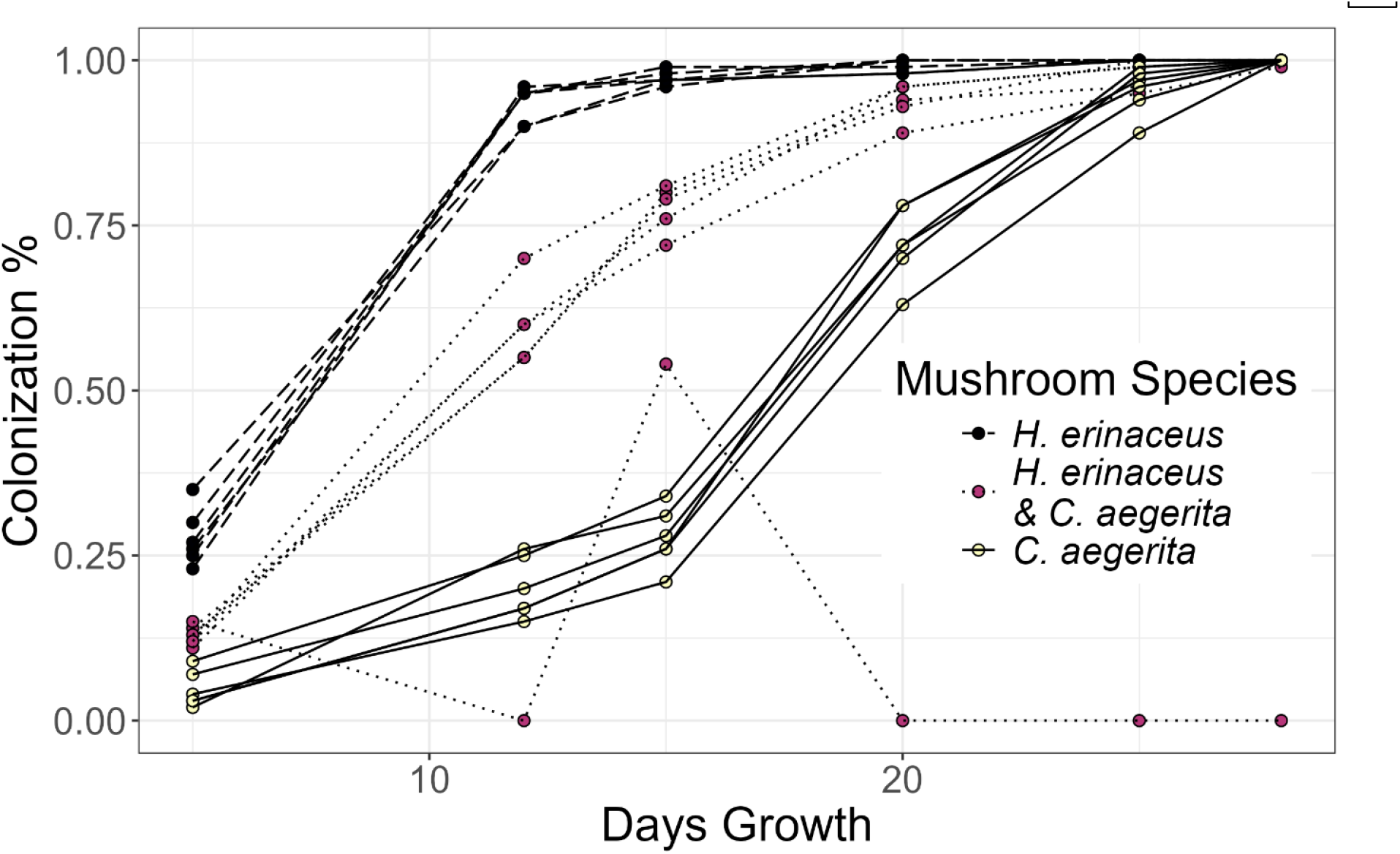
Visible hyphal colonization rate of mushroom cultivation bags with different mushrooms.

There were several contamination events in the co-cultivation and *C. aegerita* bags prepared in the facility (Fig. 2). There was a significant difference in the amount of contamination between the treatments (chisq = 29.76, DF = 2, p = 3.45e-7). When conducting pairwise comparisons between the treatments, there was a significant difference in the amount of contamination between the *H. erinaceus* and the co-cultivation bags (chisq = 31.07, DF = 1, p = 2.49e-8), with no visible contamination in the *H. erinaceus* bags at day 28. There was also a significant difference between the amount of contamination in the *C. aegerita* and co-cultivation bags (chisq = 5.87, DF = 1, p = 0.015), although several of the *C. aegerita* bags had contamination before the end of the trial. The source of the contamination was not identified, however, the logistics of inoculating a single bag with two isolates under production-scale practices could have led to contamination events.

**Figure 2.**
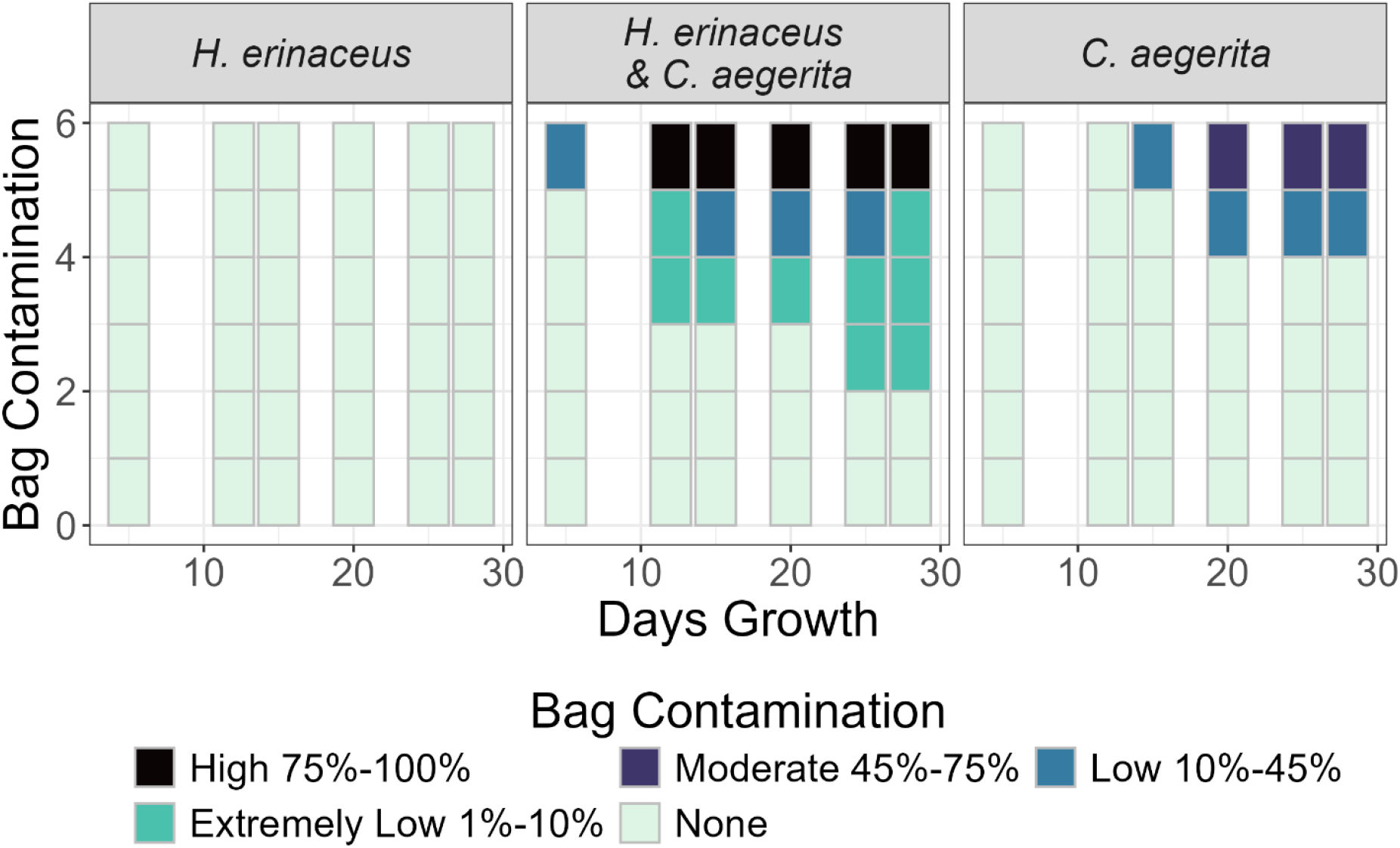
Contamination severity ratings in mushroom bags.

Several kilograms of mushrooms were produced during the experiment, in both the single isolate and co-cultivation bags (Fig. 3). The mean yield per bag of *H. erinaceus* was not significantly different when grown singly or in co-cultivation (W = 29, p = 0.094). The mean weight of mushrooms collected when grown singly was 1137.7g and in co-cultivation was 789.3g. There was a significant difference in the yield of *C. aegerita* when grown in co-cultivation, with a mean yield of 1.5g in co-cultivation and 1015.7g when grown by itself (W = 0, p = 0.0037).

**Figure 3.**
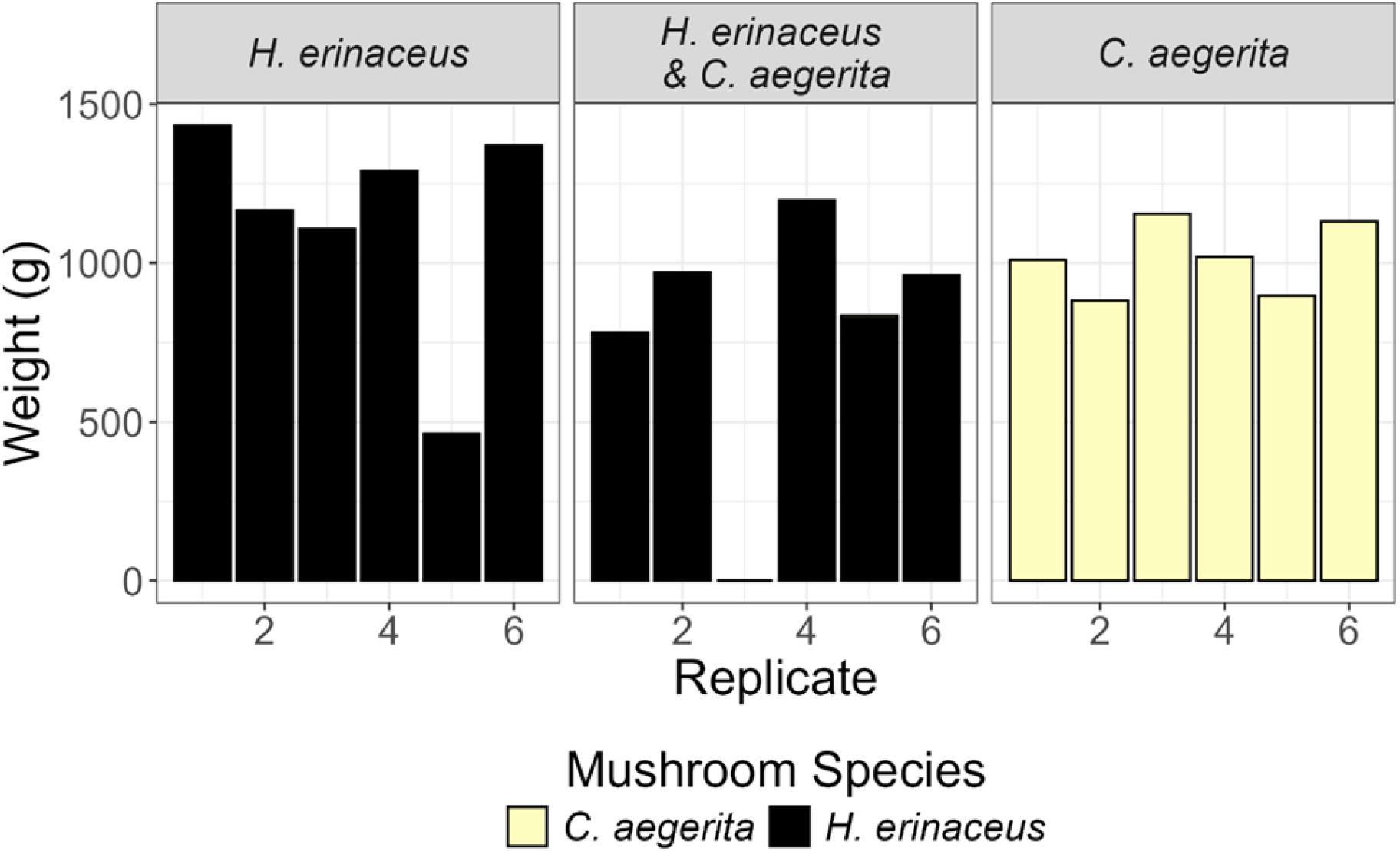
cultivated weight collected from bags of mushrooms.

## 4. Discussion

Overall, the results of co-cultivation of *C. aegerita* and *H. erinaceus* did not result in synergistic growth and yield, and instead resulted in slower overall colonization of the substrate, increased contamination, and reduced yield. The colonization rate of the media was slower with co-cultivation when compared with *H. erinaceus* alone, suggesting that some amount of antagonism between the fungi may have reduced the overall colonization rate. Relatively little yield was collected of *C. aegerita* from the co-cultivation bags, suggesting that this method may not be suitable for commercial practices. Although there was no significant reduction in yield of *H. erinaceus* when grown in co-cultivation, the increased risk of contamination offsets any potential gains that may come from the practice. Although co-cultivation methods offer promising results for increased production of novel secondary metabolites, the use of co-cultivation in production may not offer benefits for most growers.

Several features may have constrained our success in this experiment. Fungal colonization of substrates is often affected by priority effects, which dominate when one isolate arrives on the substrate prior to the other (Hiscox et al. 2015). These effects could limit the colonization of the substrate as both isolates are directly competing for resources rather than undergoing natural succession and subsequent efficient usage of spent substrate that was not fully utilized by the initial colonizers. We focused our study on *C. aegerita* and *H. erinaceus*, two white-rot fungi that utilize similar substrates(Lu et al. 2024; Rezaeian et al. 2022). It is possible that a different combination of fungal species or even different strains may result in more effective co-cultivation in commercial settings.

Optimizing mushroom production is challenging for a single species, let alone multiple species in one single growth area. It is entirely possible that the environmental conditions necessary for growth and production are different for the two species, limiting their ability to be effectively co-cultivated (Krupodorova et al. 2021). This work provides preliminary results that highlight the challenges of commercial mushroom production and the importance of research into the increasing importance and interest of mushroom production.

## Acknowledgements

This work is supported by a grant from the Education and Workforce Development Grants Program no. 2022-680183-6606 from the USDA National Institute of Food and Agriculture as well as the UTRGV Engaged Scholars and Artists Award. We would like to thank Carlos Garcia Patlan and Geovanni Hernandez for helpful feedback on experimental design and analysis. We would like to thank One Up Mushroom Production in Mission, Texas, for their invaluable resources and help.

---

**Supplemental Figure 1.**
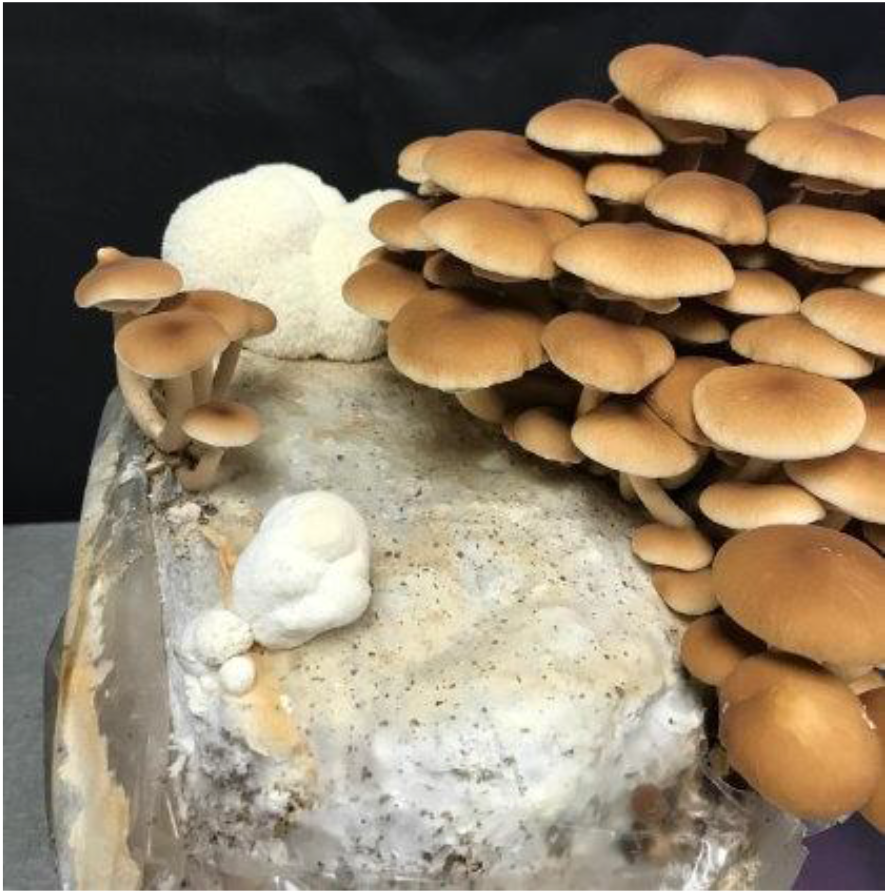
Photograph from mushroom producer showing spontaneous dual cultivation of *Cyclocybe aegerita* and *Hericium erinaceus* mushrooms on the same bag.

