## Supplementary figures and images for "Growth inhibition in co-cultivation of *Cyclocybe aegerita* and *Hericium erinaceus* mushrooms"

### Supplemental Figure 1

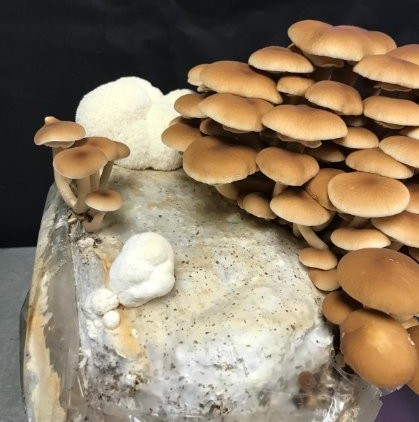
